# Ceclazepide as novel *Mycobacterium abscessus* inhibitor

**DOI:** 10.1101/2024.04.03.587875

**Authors:** K.L. Therese, J. Therese, R. Bagyalakshmi

## Abstract

The rise of drug-resistant *Mycobacterium abscessus* poses a significant challenge due to its resistance to current standard treatments, highlighting the urgent need for new antibacterial solutions. In this research, ceclazepide was introduced as a promising agent with strong activity against *M. abscessus*. Our findings revealed that ceclazepide effectively suppressed the growth of *M. abscessus* wild-type strain, multiple subspecies, and clinical isolates in vitro. Notably, ceclazepide demonstrated the ability to inhibit *M. abscessus* growth within macrophages without causing harm. These results support the potential of ceclazepide as a candidate for further development as a clinical drug to combat *M. abscessus* infections effectively.

## INTRODUCTION

Nontuberculous mycobacteria (NTM) encompass around 200 species, with the majority being environmental bacteria that do not typically cause harm to humans or animals, constituting about 95% of the group. These NTM species are commonly found in various environments frequented by humans, including surface and tap water, soil, animals, milk, and food products. Despite their benign nature, certain pathogenic NTM strains like *Mycobacterium avium* complex (MAC) and *Mycobacterium abscessus* pose significant public health concerns on a global scale (Quang and Jang, 2021). Additionally, factors such as aging, underlying pulmonary conditions like cystic fibrosis (CF), and immunocompromised states such as acquired immunodeficiency syndrome (AIDS) contribute to the heightened risk of MAC and *M. abscessus* infections in numerous regions worldwide (Johansen et al., 2020). *M. abscessus* infections pose a significant challenge, especially due to their high resistance levels (Nguyen et al., 2024). The clinical manifestations of *M. abscessus* infection are typically classified as either pulmonary or extrapulmonary disease. In particular, chronic pulmonary infections caused by *M. abscessus* are prevalent in susceptible individuals with conditions like cystic fibrosis (CF), bronchiectasis, and chronic obstructive pulmonary disease (COPD) (Wu et al., 2018; Johansen et al., 2020). Unfortunately, these infections are notoriously difficult to treat and are often considered incurable, leading to a rapid decline in lung function among affected patients. The management of *M. abscessus* infections typically involves prolonged courses of antibiotic therapy, which, regrettably, can result in high mortality rates (Wu et al., 2018; Johansen et al., 2020; Quang and Jang, 2021). Despite being less harmful to humans when compared to *M. tuberculosis*, the bacterium responsible for causing tuberculosis, treating lung infections caused by *M. abscessus* is extremely challenging. This is primarily due to the bacterium’s high resistance to commonly used antibiotics, such as β-lactams, tetracyclines, aminoglycosides, and macrolides (Luthra et al., 2018; Nguyen et al., 2024). Additionally, even anti-tuberculosis drugs like rifampicin and isoniazid prove to be ineffective against *M. abscessus*. This highlights the urgent need for innovative treatment approaches and effective strategies to combat the challenges posed by *M. abscessus* infections. There are two primary approaches to advancing the development of effective medicines targeting mycobacteria, particularly *M. abscessus* ) (Wu et al., 2018; Johansen et al., 2020). The first approach is de novo drug discovery, which involves the identification of new compounds through screening libraries of chemicals and natural products. This is followed by hit and lead development, target identification, comprehensive preclinical testing, and ultimately, clinical trials. In recent times, a number of newly developed chemical entities and antimicrobial compounds have emerged, displaying promising activity against the highly drug-resistant pathogen *M. abscessus*. Some notable examples include ohmyungsamycin, omadacycline, epetraborole, DS86760016, delpazolid, etc (Cho et al., n.d.; Kim et al., n.d.; Aziz et al., 2017; Bich Hanh et al., 2021; Egorova et al., 2021; Ganapathy et al., 2021; Jeon et al., 2022; Fujiwara et al., 2023; Nguyen et al., 2023b, 2023a). These compounds offer potential avenues for combating the challenges posed by *M. abscessus* infections. Indeed, the repurposing or repositioning of existing drugs for new therapeutic uses has shown great potential in the case of *M. abscessus*. A prime example of this is the comparison between rifampicin and rifabutin, another rifamycin analog. Rifabutin has demonstrated remarkable activity against *M. abscessus* both in vivo and in vitro (Aziz et al., 2017; Nguyen et al., 2023a). This can be attributed to its favorable oral bioavailability, excellent pharmacokinetic properties (including a long half-life and high cellular penetration), and its ability to achieve appropriate concentrations in human lung tissue. By exploring the potential of drugs already approved or in clinical use for other indications, we can uncover new treatment options for *M. abscessus* infections. These approaches hold great promise in the ongoing pursuit of improved therapies against *M. abscessus*. The 2007 American Thoracic Society NTM guidelines acknowledged the lack of a proven effective drug regimen for treating pulmonary *M. abscessus* infection . However, it did suggest a non-curative therapy involving a combination of a macrolide (clarithromycin or azithromycin) with amikacin, and cefoxitin or imipenem (Daniel-Wayman et al., 2019). More recent guidelines from the Center for Disease Control and Prevention also recommend a combination therapy with clarithromycin, amikacin, and cefoxitin as the current antimicrobial drugs of choice for treating *M. abscessus* infection (Lee et al., 2015). Nevertheless, these regimens are still subject to debate due to their ineffectiveness and significant toxic side effects over the prolonged treatment period. Therefore, *M. abscessus* poses a significant challenge in terms of antibiotic treatment, earning its reputation as an antibiotic nightmare. Consequently, there is an urgent need to discover new, effective drugs that are less toxic and more efficient in combating *M. abscessus* lung disease.

Ceclazepide, also known as TR2-A, is a novel gastrin blocker and prodrug that has demonstrated potential as an antibacterial agent against a range of bacterial pathogens (Figure 1) (Ceclazepide – Trio Medicines, n.d.; Wang et al., 2023). However, it is important to note that there have been no specific studies conducted on ceclazepide’s effectiveness against mycobacteria. The antibacterial properties of ceclazepide may be attributed to its ability to release nitric oxide, which can disrupt bacterial biofilms and hinder bacterial growth. Nevertheless, the precise mechanisms of action are still undergoing investigation (Ceclazepide – Trio Medicines, n.d.; Wang et al., 2023).

**Figure 1.**
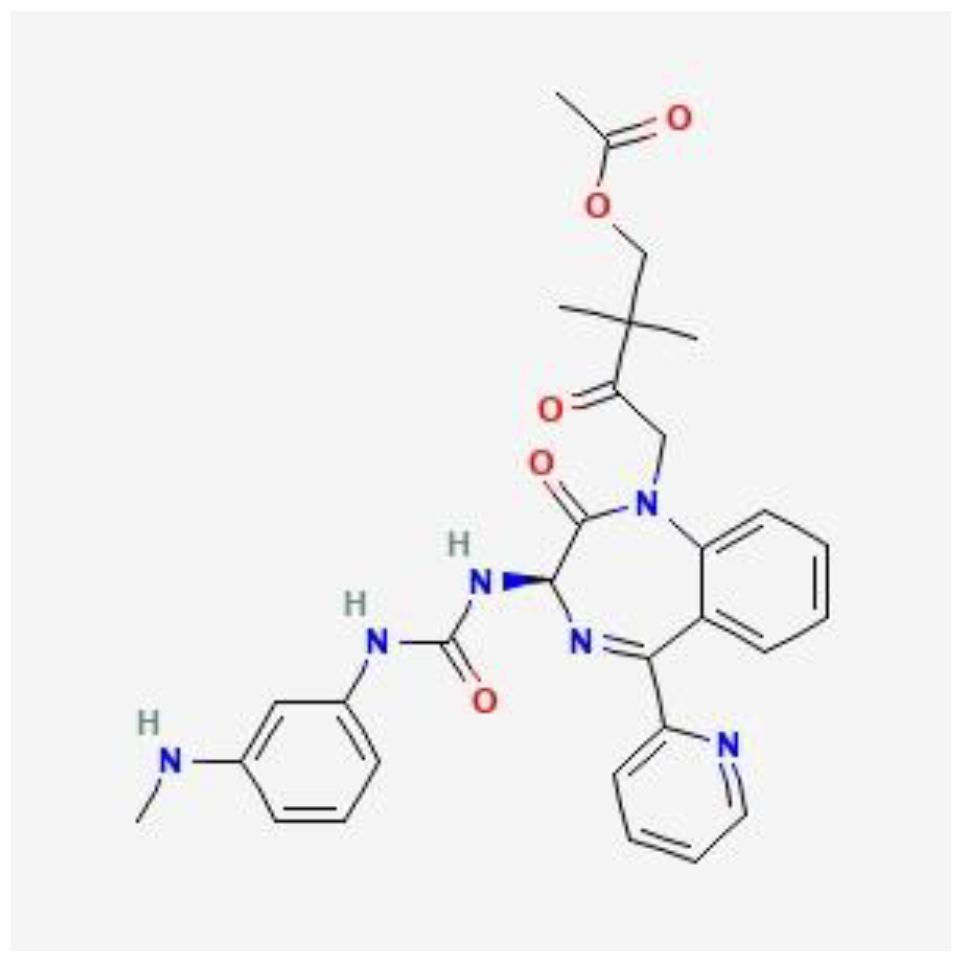
Chemical structure of ceclazepide.

In our findings, we present the discovery of a novel agent called ceclazepide, which demonstrates remarkable efficacy against *M. abscessus* both in vitro and in infected murine macrophages.

## MATERIALS AND METHODS

### Bacterial strains / culture conditions / chemicals

*M. abscessus* subsp. *abscessus* ATCC 19977 was acquired from the American Type Culture Collection, while *M. abscessus* subsp. *bolletii* CIP108541T and *M. abscessus* subsp. *massiliense* CIP108297T were obtained from the Collection de l’Institut Pasteur. The clinical isolates of *M. abscessus* were generously provided by the National University Hospital of Singapore. The *M. abscessus* strains were cultivated at a temperature of 37°C in a Middlebrook 7H9 culture medium (Difco), which was supplemented with 10% albumin-dextrose-catalase (ADC, Difco) and 0.05% Tween-80 (Sigma). For determining the colony-forming units (CFU), the bacteria were plated on a solid culture medium called Middlebrook 7H10, which contained 0.5% glycerol and 10% OADC (Difco). To assess the minimum inhibitory concentration (MIC), the bacteria were tested in a cation-adjusted Mueller-Hinton (CAMH) medium (Sigma, St. Louis, MO, USA), supplemented with 20 mg/L calcium chloride (Sigma, St. Louis, MO, USA) and 10 mg/L magnesium chloride (Sigma, St. Louis, MO, USA). All cultures were incubated at 37°C with continuous shaking at 180 rpm. The ceclazepide and clarithromycin used in the experiments were purchased from DC Chemicals (Shanghai, China).

### Resazurin Microtiter Assay (REMA)

The determination of MIC values was carried out using the resazurin microtiter assay (REMA). In this assay, bacterial stocks from the exponential-phase cultures were diluted to an optical density at 600 nm (OD600) of 0.01. Each well of a sterile, polystyrene 96-well cell culture plate was then filled with 50 μL of the bacterial culture, and 50 μL of serial 2-fold dilutions of the test compound solution were added to each well. A drug-free growth control was included on each plate. To prevent evaporation during incubation, 200 μL of sterile water was added to the outer perimeter wells. The plates were covered with self-adhesive membranes and incubated at 37°C for 3 days. Afterward, 40 μL of a 0.025% (wt/vol) resazurin solution was added to each well, and the plates were reincubated overnight. The fluorescence was measured using a SpectraMax M3 multimode microplate reader (Molecular Devices, Sunnyvale, CA, USA). A dose-response curve was constructed, and the concentrations required to inhibit bacterial growth by 50% (MIC50) were determined using GraphPad Prism software (version 6.05; San Diego, CA, USA).

### Isolation of BMDMs and intracellular bacterial replication assay

Bone marrow-derived macrophages (BMDMs) were obtained by flushing the femur and tibia of 6-week-old C57BL/6 mice. The BMDMs were cultured in Dulbecco’s modified Eagle’s medium (DMEM; Welgene, Gyeongsan-si, Gyeongsangbuk-do, South Korea) supplemented with 10% fetal bovine serum (FBS; Welgene), GlutaMax (35050-061; Gibco), and penicillin/streptomycin (15140-122; Gibco) at 37°C with 5% CO2. To differentiate the cells into mature BMDMs, they were exposed to recombinant murine macrophage colony-stimulating factor (M-CSF; JW-M003-0025, JW CreaGene) for a period of 5 days.

To test the antibacterial activity of ceclazepide against intracellular bacteria, the BMDMs were infected with *M. abscessus* subsp. *abscessus* ATCC 19977 at a multiplicity of infection of 1 in 96-well plates for a duration of 3 hours. After the infection, extracellular bacteria were eliminated by treating the cells with 250 μg/mL amikacin for 2 hours. The cells were then washed with phosphate-buffered saline (PBS; Gibco) and treated with serially diluted compounds in 96-well plates for a period of 3 days. To release the intracellular bacteria, the cells were lysed using 1% SDS (151-21-3; Generay Biotechnology). The lysates were subsequently 10-fold serially diluted with PBS, and each dilution was plated on 7H10-OADC supplemented with 50 μg/mL kanamycin. The plates were incubated for at least 3 days at 37°C, after which the bacterial colonies were counted.

## RESULTS

### Ceclazepide Exhibits Potent Activity against M. abscesuss Subspecies and Clinical Isolates

To assess the activity of ceclazepide, we determined the minimum concentrations required to inhibit 50% of the growth of different *M. abscessus* subspecies, including *M. abscessus* subsp. *abscessus* ATCC 19977 smooth (S) and rough (R) morphotypes, *M. abscessus* subsp. *massiliense* CIP108297T, and *M. abscessus* subsp. *bolletii* CIP108541T. Clarithromycin was used as a positive control. After incubating the *M. abscessus* strains with the compounds for 3 days, a dose-dependent decrease in fluorescence was observed, indicating a killing effect. As shown in Figure 2, all subspecies tested were susceptible to ceclazepide. The MIC50 values of ceclazepide ranged from 1.8 to 8.2 μM (MIC90 values ranged from 4.3 to 28.3 μM). Notably, *M. abscessus* subsp. *massiliense* CIP108297T exhibited the lowest MIC50 value of 4.4 μM (MIC90 value of 13.7 μM), while *M. abscessus* subsp. *abscessus* ATCC 19977 showed the highest MIC50 value of 8.2 μM (MIC90 value of 28.4 μM). *M. abscessus* subsp. *bolletii* CIP108541T had a MIC50 value of 6.1 μM (MIC90 value of 7.1 μM). Furthermore, we evaluated the activity of ceclazepide against clinical isolates of *M. abscessus*. As shown in Figure 2, significant growth inhibition was observed when the clinical isolates were treated with various concentrations of ceclazepide. The MIC50 values ranged from 7.8 to 9.9 μM (MIC90 values ranged from 7.1 to 28.3 μM). Clarithromycin, used as a positive control, also inhibited the growth of *M. abscessus* subsp. *abscessu*s ATCC 19977 with a similar MIC50 value to ceclazepide for all tested *M. abscessus* strains. Based on these results, ceclazepide can be considered an effective drug candidate for *M. abscessus* strains.

**Figure 2.**
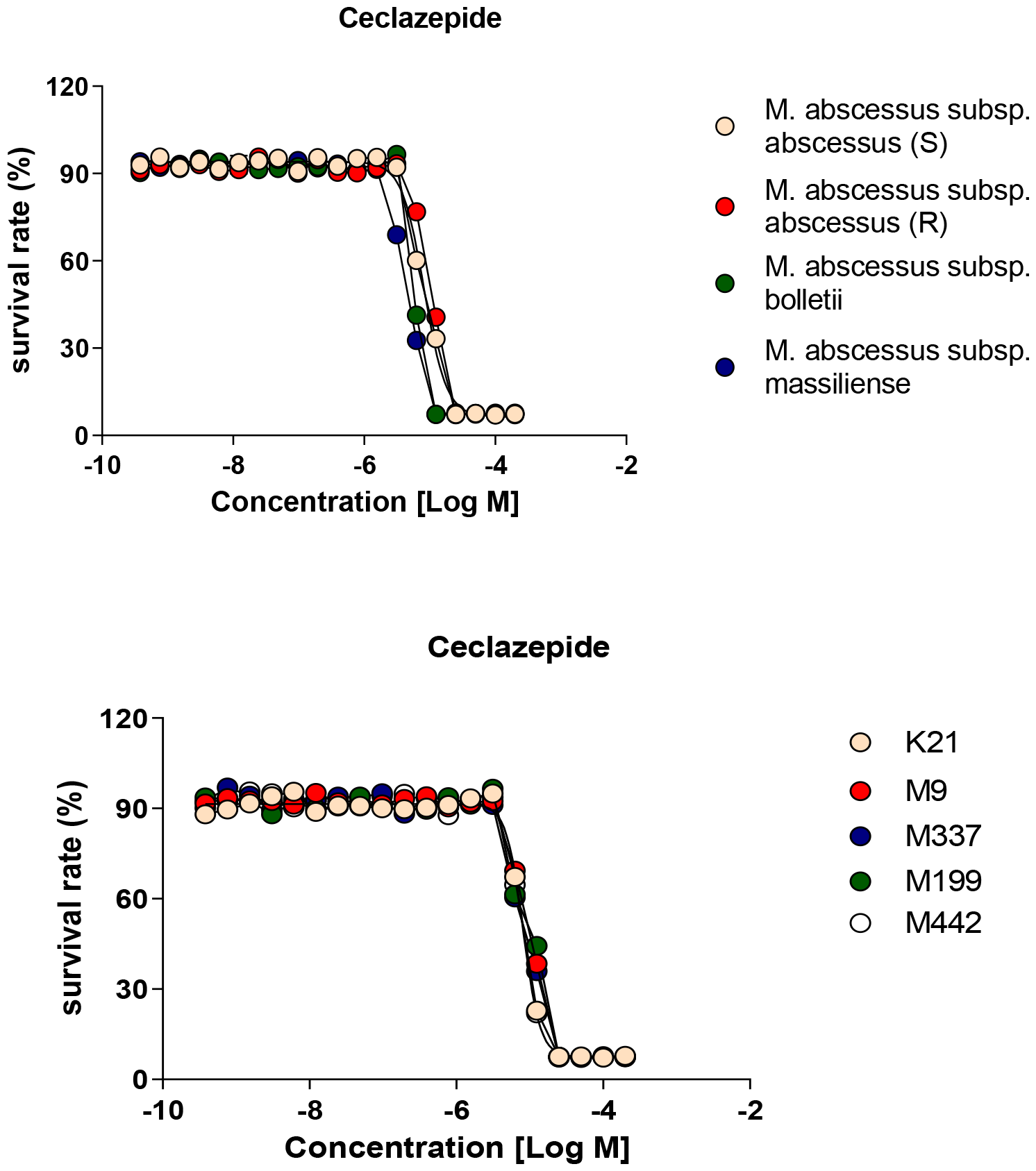
In vitro activity of ceclazepide against *M. abscessus* strains.

### Ceclazepide activity against intracellular replicating M. abscessus

To assess the efficacy of ceclazepide against *M. abscessus* replication within host cells, mouse bone marrow-derived macrophages (mBMDMs) were infected with *M. abscessus* subsp. *abscessus* ATCC 19977. Clarithromycin and dimethyl sulfoxide (DMSO) were used as the positive and negative controls, respectively. The number of viable *M. abscessus* subsp. *abscessus* ATCC 19977 inside the mBMDMs was determined using the traditional colony count method after lysing the infected cells treated with ceclazepide and clarithromycin, respectively. The results of the study showed that ceclazepide treatment resulted in a significant decrease in the number of intracellular mycobacteria following infection. Ceclazepide exhibited a strong reduction in *M. abscessus* which was comparable to the reduction observed with clarithromycin with IC50 of 1.3 and 2.5 μM respectively (Figure 3). Based on these findings, it can be concluded that ceclazepide demonstrated activity against intracellular *M. abscessus*.

**Figure 3.**
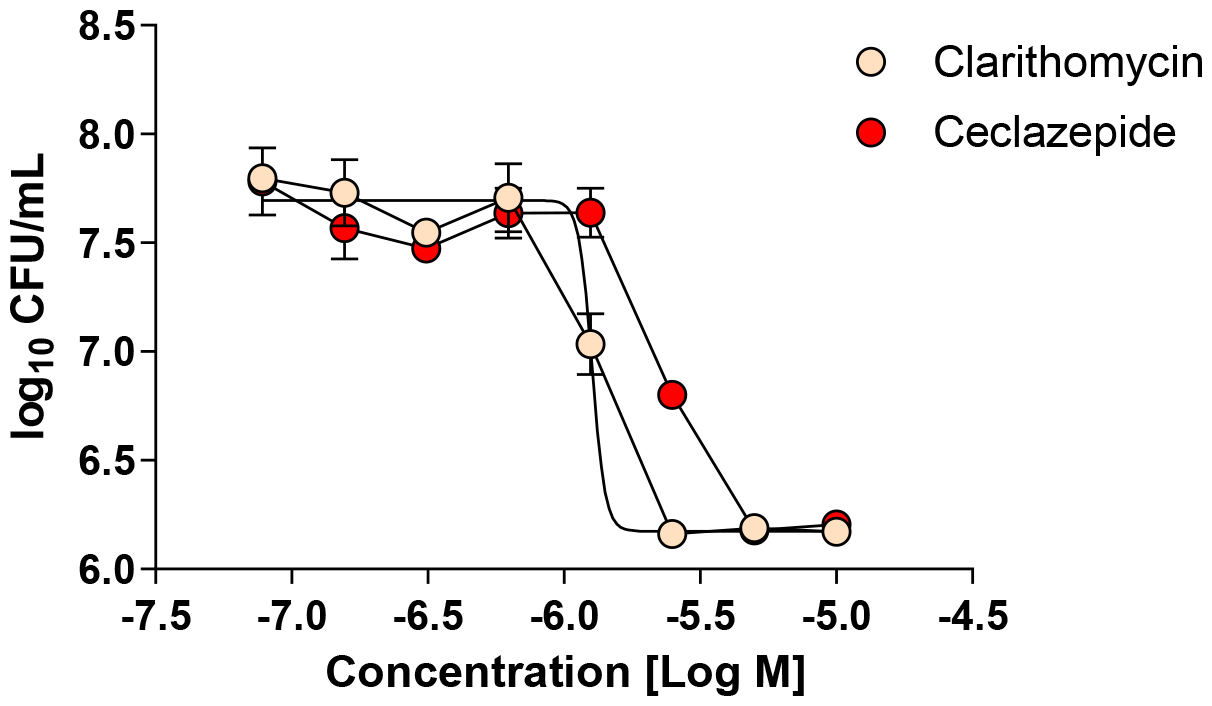
Intracellular activity of ceclazepide against *M. abscessus*.

### DISCUSSION

*M. abscessus* is a highly drug-resistant mycobacterial species, and the lack of effective treatment options is a significant challenge (Luthra et al., 2018; Johansen et al., 2020; Quang and Jang, 2021). The current clarithromycin-based treatment regimens have only moderate efficacy, and the emergence of clarithromycin resistance in some isolates leads to treatment failure. Therefore, the discovery of new chemical entities with potent activity against *M. abscessus* is of utmost importance. Various approaches, such as whole-cell screening, combination studies with existing drugs to achieve synergistic effects, and drug repositioning studies with known compounds, have been conducted in recent years. However, there is still a lack of promising new chemical leads that have progressed to clinical phase III and market release. Currently, only three clinical phase II studies have been completed to evaluate the safety, tolerance, and efficacy of potential treatments. These include tigecycline, an inhaled formulation of nitric oxide (NO), and a novel formulation of liposomal amikacin for inhalation (LAI). However, no results from these trials have been posted on ClinicalTrials.gov as of yet. Furthermore, there are limited active new drug candidates in clinical phase I studies, even at the discovery level. The low hit rate in chemical drug screens targeting *M. abscessus* may contribute to this scarcity (Wu et al., 2018). Therefore, there is an urgent need to identify active agents for *M. abscessus*, despite the fact that the minimum inhibi tory concentration (MIC) values are in the micromolar range.

Ceclazepide, also known as TR2-A, is a novel gastrin blocker and prodrug that has shown promise as an antibacterial agent against various bacterial pathogens (Ceclazepide – Trio Medicines, n.d.; Wang et al., 2023). While there have been no specific studies conducted on ceclazepide’s effectiveness against mycobacteria, its antibacterial properties may be linked to its ability to release nitric oxide (Ceclazepide – Trio Medicines, n.d.; Wang et al., 2023). Nitric oxide has been shown to disrupt bacterial biofilms and inhibit bacterial growth (Ceclazepide – Trio Medicines, n.d.; Wang et al., 2023). However, further research is needed to fully understand the mechanisms of action of ceclazepide. In this study, we conducted three different evaluations to assess the activity of ceclazepide, and the results consistently demonstrated its effectiveness against *M. abscessus*. Firstly, we examined the in vitro susceptibility of *M. abscessus* to ceclazepide. The presence of ceclazepide significantly reduced the survival of all three *M. abscessus* subspecies, and its in vitro activity was comparable to that of clarithromycin. Additionally, we investigated the antibacterial activity of ceclazepide against clinical strains of *M. abscessus* and found that it exhibited equal efficacy across different *M. abscessus* clinical isolates. These findings highlight the potential of ceclazepide as an effective treatment option for *M. abscessus* infections.

The study aimed to evaluate the intracellular antimicrobial activity of ceclazepide against *M. abscessus*, a bacterium that replicates inside murine bone marrow-derived macrophages (mBMDMs). During infection, *M. abscessus* evades host defenses by being phagocytized by macrophages and replicating within immune cells. Therefore, identifying compounds that can effectively kill intracellular *M. abscessus* by penetrating the cell membrane is crucial. The results of the study demonstrated that ceclazepide treatment led to a significant decrease in the number of intracellular mycobacteria following infection. Ceclazepide exhibited a reduction in *M. abscessus* comparable to the reduction observed with clarithromycin. These findings indicate that ceclazepide possesses activity against intracellular *M. abscessus*.

The results obtained strongly support the potential of ceclazepide as a promising candidate for further development as a clinical drug to effectively combat *M. abscessus* infections. These findings highlight the efficacy of ceclazepide and suggest its potential as a valuable addition to the armamentarium against *M. abscessus*. Further research and clinical trials are warranted to explore its full therapeutic potential and assess its safety and efficacy in treating *M. abscessus* infections.

## AUTHOR CONTRIBUTIONS

RB concptualised the research work. KL, JT, and RB searched and gathered the previous studies. RB wrote the manuscript. KL, JT, and RB critically reviewed the manuscript. RB edited the reviewed manuscript. RB critically evaluated and revised the manuscript and supervised the whole project.

## ACKNOWLEDGMENTS

The authors would like to acknowledge the Indian Council of Medical Research (ICMR), New Delhi, for their financial support in conducting this study. Their support has been instrumental in enabling the research and contributing to the findings presented in this work.

