## supplemental for "Ceclazepide as novel *Mycobacterium abscessus* inhibitor"

### Slide 1
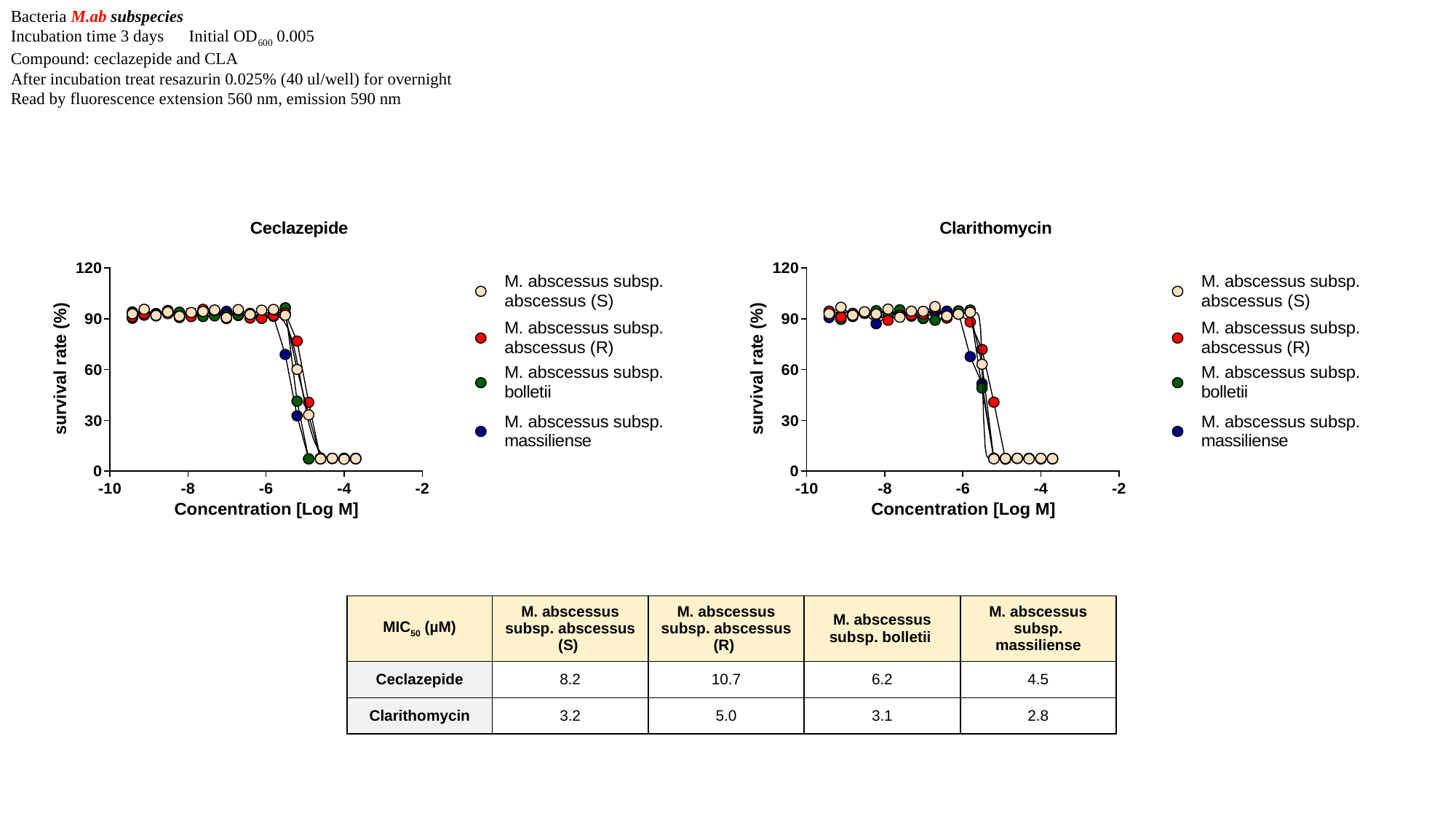

Bacteria M.ab subspecies
Incubation time 3 days Initial OD_600_ 0.005
Compound: ceclazepide and CLA
After incubation treat resazurin 0.025% (40 ul/well) for overnight
Read by fluorescence extension 560 nm, emission 590 nm
| MIC_50_ (µM) | M. abscessus subsp. abscessus (S) | M. abscessus subsp. abscessus (R) | M. abscessus subsp. bolletii | M. abscessus subsp. massiliense |
| --- | --- | --- | --- | --- |
| Ceclazepide | 8.2 | 10.7 | 6.2 | 4.5 |
| Clarithomycin | 3.2 | 5.0 | 3.1 | 2.8 |

### Slide 2
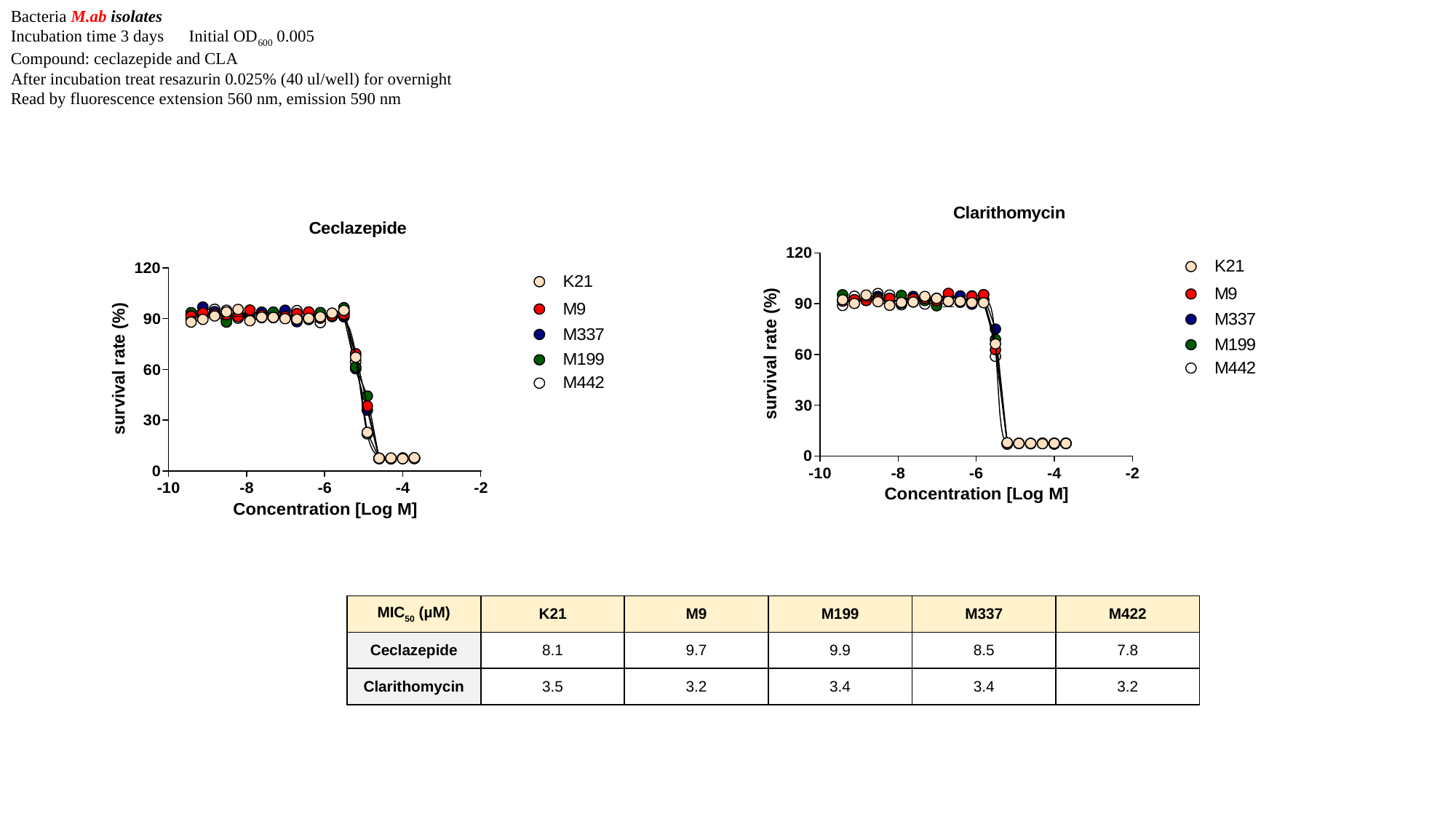

Bacteria M.ab isolates
Incubation time 3 days Initial OD_600_ 0.005
Compound: ceclazepide and CLA
After incubation treat resazurin 0.025% (40 ul/well) for overnight
Read by fluorescence extension 560 nm, emission 590 nm
| MIC_50_ (µM) | K21 | M9 | M199 | M337 | M422 |
| --- | --- | --- | --- | --- | --- |
| Ceclazepide | 8.1 | 9.7 | 9.9 | 8.5 | 7.8 |
| Clarithomycin | 3.5 | 3.2 | 3.4 | 3.4 | 3.2 |

### Slide 3
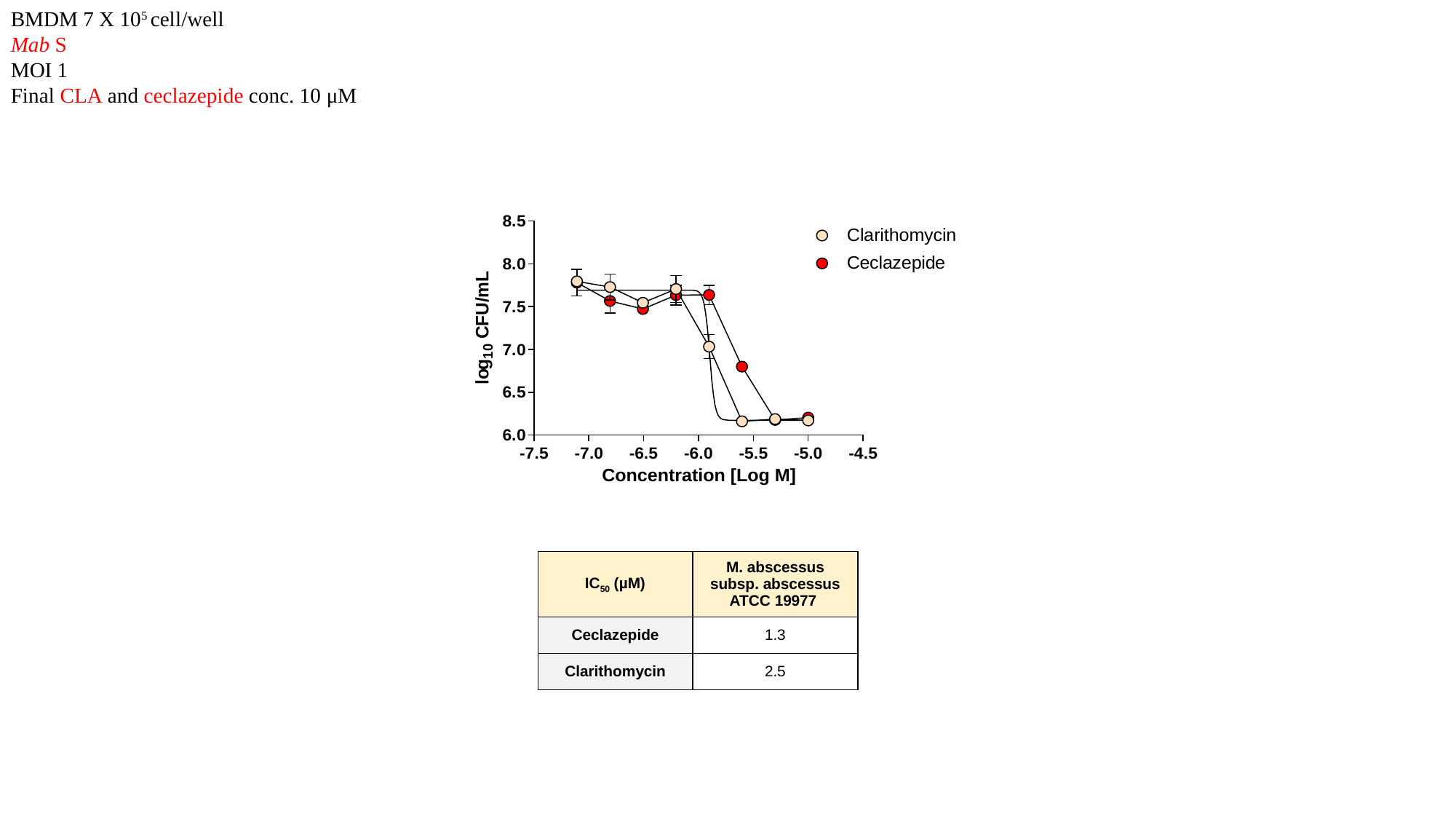

BMDM 7 X 10^5^ cell/well
Mab S
MOI 1
Final CLA and ceclazepide conc. 10 μM
| IC_50_ (µM) | M. abscessus subsp. abscessus ATCC 19977 |
| --- | --- |
| Ceclazepide | 1.3 |
| Clarithomycin | 2.5 |
